## Supplemental Tables 1-4 for "Mycobacterial resistance to zinc poisoning requires assembly of P-ATPase-containing membrane metal efflux platforms"

**SUPPLEMENTARY TABLES S1-S4**

Table S1. Related to Materials and Methods; Primers used for the construction of Mycobacterial mutant strains.

| ***Mycobacterium tuberculosis* strains** | |
| --- | --- |
| **Construction of H37Rv Δ(*pacL1-ctpC*)::Zeo^R^** | |
| *PCR amplification of pacL1 upstream fragment* | |
| 3269-Am-Fw | AGCGATACTCGACGATTC |
| 3269-Am-Rv-Zeo | CAGTCGATCCACGTGGAGCATTGCCTGTACCTTTCTTCC |
| *PCR amplification of ctpC downstream fragment* | |
| 3270-Zeo-Av-Fw | CCACTGAGCGTCAGACCCACGTGCTCGCCAACAGTTCCCGGTTGATCC |
| 3270-Av-Rev | AGCCTGGCGGTATTGCTCAC |
| **Construction of H37Rv Δ*pacL1*::*dif4*** | |
| *PCR amplification of pacL1 upstream fragment* | |
| 3269-Am-Fw | AGCGATACTCGACGATTC |
| 3269-Am-Rv-Zeo2 | CAGTCGATCCACGTGGAGGCCATTGCCTGTACCTTTCTTCC |
| *PCR amplification of pacL1 downstream fragment* | |
| 3269-Zeo-Av-Fw | CCACTGAGCGTCAGACCCACGTGCTCACCTTGAACTCGCCAGGAC |
| Seq-ctpC-Rev1 | TTCTCGCGCAGGATGAGGCTTG |
| **Construction of H37Rv ΔRv1488::*dif6*** | |
| *PCR amplification of Rv1488 upstream fragment* | |
| Rv1488-Am-Fw | GGCTGACGCGGATCACGCG |
| Rv1488-Am-Rv-Zeo | CAGTCGATCCACGTGGAGACGGCTCCTTGCACCGGAATTCCTTTC |
| *PCR amplification of Rv1488 downstream fragment* | |
| Rv1488-Zeo-Av-Fw | CCACTGAGCGTCAGACCCACGTGCTCTCAATAGAGTGGTCCGATGAGTGG |
| Rv1488-Av-Rv | CTAGTGCGGCGTCGACCC |
| **Construction of H37Rv Δ(*pacL1::dif4 pacL2::dif6 pacL3::dif5*)** | |
| *PCR amplification of pacL2 upstream fragment* | |
| Rv1993c-Am-Fw | CGTGAGTTGGCGGCTCTG |
| Rv1993c-Am-Rv-Zeo | CAGTCGATCCACGTGGAGGTAACCACGGTCAGTTCTCC |
| *PCR amplification of pacL2 downstream fragment* | |
| Rv1993c-Zeo-Av-Fw | CCACTGAGCGTCAGACCCACGTGCTCCGAGTGACGACTGTAGTTGACGCCGAG |
| Rv1993c-Av-Rv | CAGCGGATAAGCCCATGCGG |
| *PCR amplification of pacL3 upstream fragment* | |
| Rv0968-Up-Fw | GTTGCTCATCGTCCACGAG |
| Rv0968-Up-Rv-Zeo | CAGTCGATCCACGTGGAGCCATGCCACACCATCGCGGATG |
| *PCR amplification of pacL3 downstream fragment* | |
| Rv0968-Zeo-Dw-Fw | CCACTGAGCGTCAGACCCACGTGCTCCCACTGACGTTCTTTCTGACACCG |
| Rv0968-Dw-Rv | GGTAAGCACCGAAGAACATTG |
| ***Mycobacterium smegmatis* strains** | |
| **Construction of mc²155 Δ*(MSMEG_0755)*::*dif4*** | |
| *PCR amplification of MSMEG_0755 upstream fragment* | |
| SMEG0755-Am-Fw | CAGGCGTGCGATTCCGTGCG |
| SMEG0755-Am-Rv-Zeo | CAGTCGATCCACGTGGAGGTGGCCCGCGCCCATTCC |
| *PCR amplification of MSMEG_0755 downstream fragment* | |
| SMEG0755-Zeo-Av-Fw | CCACTGAGCGTCAGACCCACGTGCTCGCAGGTGCGAGGCCGAGTGAC |
| SMEG0755-Av-Rv | CGATGGATTCGGCATCCCTTTTGCG |
| **Construction of mc²155 Δ*(MSMEG_6059-6058)*::*dif5*** | |
| *PCR amplification of MSMEG_6059 upstream fragment* | |
| SMEG6059-Am-Fw | GCAAGCATGAACGTCAATGCG |
| SMEG6059-Am-Rv-Zeo | CAGTCGATCCACGTGGAGCCATGCCACACCATCGGCCAAGC |
| *PCR amplification of MSMEG_6058 downstream fragment* | |
| SMEG6058-Zeo-Av-Fw | CCACTGAGCGTCAGACCCACGTGCTCCGACTAGGGCGTTCGGCGAACG |
| SMEG6058-Av-Rv | GTGAGTGGTTTCACTCCG |

**Table S2. Related to Materials and Methods; Plasmids used in this study and primers used for their construction.**

| **Plasmid** | **Parent vector/ Selection** | **Cloning technique** | **Plasmids, Primer pairs or Restriction enzymes used** | **Reference** |
| --- | --- | --- | --- | --- |
| pJV53H | pJV53; hygro^R^ | - | - | ^1^ |
| pDE43-MCS | Strepto^R^ | - | Destination vector for multisite gateway cloning | ^2^ |
| pDO41A | Ampi^R^ | - | Destination vector to construct pEN41 5’ entry plasmids for gateway cloning | ^2^ |
| pDO12A | Ampi^R^ | - | Destination vector to construct pEN12 middle entry plasmids for gateway cloning | ^2^ |
| pDO23A | Ampi^R^ | - | Destination vector to construct pEN23 3’ entry plasmids for gateway cloning | ^2^ |
| pEN41A-T02 | Ampi^R^ | - | Empty 5’ entry clone for multisite gateway cloning | ^2^ |
| pEN12A-AgeI | Ampi^R^ | - | Empty middle entry clone for multisite gateway cloning | ^2^ |
| pEN23A-MluI | Ampi^R^ | - | Empty 3’ entry clone for multisite gateway cloning | ^2^ |
| pEN23A-*P_pacL1_*-*pacL1* | pDO23A Ampi^R^ | Gateway cloning | PCR using H37Rv chromosomal DNA as template and primers clo-B2-3269-Am/clo-B3-3269-Av | This work |
| pEN23A-*P_pacL1_*-*pacL1-ctpC* | pDO23A Ampi^R^ | Gateway cloning | PCR using H37Rv chromosomal DNA as template and primers clo-B2-3269-Am/clo-B3-ctpC-Av | This work |
| pEN41A-P1-mCherry | pDO41A Ampi^R^ |  | Donor of constitutively expressed mCherry reporter for multisite gateway | ^3^ |
| pLAM12::GFP | - | - | Source of GFP sequence | ^3^ |
| pEN23A-*P_pacL1_*-GFP | pDO23A Ampi^R^ | Gateway cloning | B2-PRv3269-Fw/PRv3269-GFP-Rev to amplify a *P_pacL1_* DNA fragment and PRv3269-GFP-Fw/B3-GFP-Rev to amplify a GFP DNA fragment. A fused fragment amplified by two-fragment PCR with B2-PRv3269-Fw/B3-GFP-Rev primers | This work |
| pGMCS-P1-mCherry | pDE43-MCS Strepto^R^ | Gateway cloning | pEN41A-P1-mCherry + pEN12A-AgeI + pEN23A-MluI + pDE43-MCS | This work |
| pGMCS-P1-mCherry-*P_PacL1_*-GFP | pDE43-MCS Strepto^R^ | Gateway cloning | pEN41A-P1-mCherry + pEN12A-AgeI + pEN23A-*P_pacL1_*-GFP + pDE43-MCS | This work |
| pGMCS-*P_pacL1_-pacL1* | pDE43-MCS Strepto^R^ | Gateway cloning | pEN41A-T02 + pEN12A-AgeI + pEN23A-*P_pacL1_*-*pacL1* + pDE43-MCS | This work |
| pGMCS-*P_pacL1_-pacL1-ctpC* | pDE43-MCS Strepto^R^ | Gateway cloning | pEN41A-T02 + pEN12A-AgeI + pEN23A-*P_pacL1_*-*pacL1-ctpC* + pDE43-MCS | This work |
| pGMCS-*P_pacL1_-ctpC* | pDE43-MCS Strepto^R^ | In-fusion cloning | PCR using pGMCS-*P_pacL1_-pacL1-ctpC* as template with pGMC 3269 Am Left/pGMC del3269 Right primers and circularization by In-fusion reaction | This work |
| pGMCS-*P_pacL1_-PacL1*Δ3 | pDE43-MCS Strepto^R^ | In-fusion cloning | PCR using pGMCS-*P_pacL1_-pacL1* as template with Del3269-Cterm-Left/Del3269-Cterm3-Right primers and circularization by In-fusion reaction | This work |
| pGMCS-*P_pacL1_-PacL1*Δ7 | pDE43-MCS Strepto^R^ | In-fusion cloning | PCR using pGMCS-*P_pacL1_-pacL1* as template with Del3269-Cterm-Left/Del3269-Cterm7-Right primers and circularization by In-fusion reaction | This work |
| pGMCS-*P_pacL1_-pacL1*(ΔTM) | pDE43-MCS Strepto^R^ | In-fusion cloning | PCR using pGMCS-*P_pacL1_-pacL1* as template with Del3269-Nterm-Left/Del3269-Nterm7_26-Right primers and circularization by In-fusion reaction | This work |
| pGMCS-*P_pacL1_*-*pacL1*_Flag_-*ctpC* | pDE43-MCS Strepto^R^ | In-fusion cloning | PCR using pGMCS-*P_pacL1_-pacL1-ctpC* as template with pGMC-Cter-rv3269 Left #3/3269 Cter Flag-Am-ctpC-Right primers and circularization by In-fusion reaction | This work |
| pGMCS-*P_pacL1_*-*pacL1*_Flag_-*ctpC*_His6_ | pDE43-MCS Strepto^R^ | In-fusion cloning | PCR using pGMCS-*P_pacL1_-pacL1*_Flag_*-ctpC* as template with pGMC-Cter-rv3269 Left #1/ctpC Cter His-Av-attB3-Right primers and circularization by In-fusion reaction | This work |
| pGMCS-*P_pacL1_*-*pacL1*(Δ7)_Flag_-*ctpC*_His6_ | pDE43-MCS Strepto^R^ | In-fusion cloning | PCR using pGMCS-*P_pacL1_-pacL1*_Flag_*-ctp C*_His6_ as template with pGMC-Cter-ctpC-left #1/ctpC Cter His-Av-attB3-Right primers and circularization by In-fusion reaction | This work |
| pGMCS-*P_pacL1_*-*pacL1*(ΔTM)_Flag_-*ctpC* | pDE43-MCS Strepto^R^ | In-fusion cloning | PCR using pGMCS-*P_pacL1_-pacL1*_Flag_*-ctp C* as template with Del3269-Nterm-Left/Del3269-Nterm7_26-Right primers and circularization by In-fusion reaction | This work |
| pJYB234 | Cam^R^ | - | Source of mVenus sequence | ^4^ |
| pJYB240 | Cam^R^ | - | Source of mTurquoise sequence | ^5^ |
| pGMCS-*P_pacL1_*-*pacL1*_mT_-*ctpC* | pDE43-MCS Strepto^R^ | In-fusion cloning | m-Turquoise coding fragment amplified with pJYB240 template and mTurquoise-Fw #3/mTurquoise-Rv#3 primers cloned by In-fusion reaction in the backbone amplified from pGMCS-*P_pacL1_*-*pacL1*-*ctpC* with pGMC-Cter-rv3269 Left #3/pGMC-Am-ctpC Right #3 | This work |
| pGMCS-*P_pacL1_*(ΔPB)-*pacL1*_mT_-*ctpC* | pDE43-MCS Strepto^R^ | In-fusion cloning | PCR using pGMCS-*P_pacL1_*-*pacL1*_mT_-*ctpC* with Inf-ctpC-MutPB-L1/Inf-ctpC-MutPB-R1 primers and circularization by by In-fusion reaction | This work |
| pGMCS-*P_pacL1_*-*pacL1*-*ctpC*_mV_ | pDE43-MCS Strepto^R^ | In-fusion cloning | m-Venus coding fragment amplified with pJYB234 template and mVenus-Fw #1/mVenus-Rv#1 primers cloned by In-fusion reaction in the backbone amplified from pGMCS-*P_pacL1_*-*pacL1*-*ctpC* with pGMC-Cter-ctpC Left #1/pGMC-Av-attB3-Right #1 | This work |
| pGMCS-*P_pacL1_*-*pacL1*_mT_-*ctpC*_mV_ | pDE43-MCS Strepto^R^ | In-fusion cloning | m-Turquoise coding fragment amplified with pJYB240 template and mTurquoise-Fw #3/mTurquoise-Rv#3 primers cloned by In-fusion reaction in the backbone amplified from pGMCS-*P_pacL1_*-*pacL1*-*ctpC*_mV_ with pGMC-Cter-rv3269 Left #3/pGMC-Am-ctpC Right #3 | This work |
| pGMCS-*P_pacL1_*-pacL1(ΔTM)_mT_-*ctpC* | pDE43-MCS Strepto^R^ | In-fusion cloning | PCR using pGMCS-*P_pacL1_*-*pacL1*_mT_-*ctpC* as template with Del3269-Nterm-Left/Del3269-Nterm7_26-Right primers and circularization by In-fusion reaction | This work |
| pGMCS-*P_pacL1_*-*pacL1*_mT_-Rv1488_mV_ | pDE43-MCS Strepto^R^ | In-fusion cloning | Rv1488 coding fragment amplified with H37Rv genomic DNA and infus Rv1488 Fw/infus mVenus Rv1488 Rv primers cloned by In-fusion reaction in the backbone amplified from pGMCS-*P_pacL1_*-*pacL1*_mT_-*ctpC*_mV_ with primers mT av left/infus backbone mVenus Right | This work |
| pEN41A-P1-mCherry | pDO41A Ampi^R^ | - | - | ^3^ |
| pEN41A-P1-Rv1488_mC_ | pDO41A Ampi^R^ | In-fusion cloning | Rv1488 coding fragment amplified from pGMCS-*P_pacL1_*-*pacL1*_mT_-Rv1488_mV_ and infus P1 Rv1488 Fw/pGMC-3269-FLUO-ctpC Left primers cloned by In-fusion reaction into the backbone amplified from pEN41A-P1-mCherry with mCherry Fw2/Infus pEN41A-P1_x_mCherry Right primers | This work |
| pGMCS-P1-Rv1488_mC_-*P_pacL1_*-*pacL1*_mT_-*ctpC*_mV_ | pDE43-MCS Strepto^R^ | In-fusion cloning | Rv1488_mC_ fragment amplified with pEN41A-P1-Rv1488_mC_ and infus P1 Rv1488 Fw (pGMC)/infus mCherry Rv cloned by In-fusion reaction in the backbone amplified from pGMCS-*P_pacL1_*-*pacL1*_mT_-*ctpC*_mV_ with pGMC-12-left/pGMC-12-right | This work |
| pGMCS-*P_pacL1_*-*pacL1*(Δ37-86)_Flag_-*ctpC*_His6_ | pDE43-MCS Strepto^R^ | In-fusion cloning | PCR using pGMCS-*P_pacL1_*-*pacL1*_Flag_-*ctpC*_His6_ as template with Inf-3269Del(37-86)-left/Inf-3269C7-right primers and circularization by In-fusion reaction | This work |
| pGMCS-*P_pacL1_*-*pacL1*(Δ54-86)_Flag_-*ctpC*_His6_ | pDE43-MCS Strepto^R^ | In-fusion cloning | PCR using pGMCS-*P_pacL1_*-*pacL1*_Flag_-*ctpC*_His6_ as template with Inf-3269Del(53-86)-left/Inf-3269C7-right primers and circularization by In-fusion reaction | This work |
| pGMCS-*P_pacL1_*-*pacL1*(Δ37-86)_mT_-*ctpC*_mV_ | pDE43-MCS Strepto^R^ | In-fusion cloning | PCR using pGMCS-*P_pacL1_*-*pacL1*_mT_-*ctpC*_mV_ as template with Inf-3269Del(37-86)-left/Inf-3269C7-right primers and circularization by In-fusion reaction | This work |
| pGMCS-*P_pacL1_*-*pacL1*(Δ54-86)_mT_-*ctpC*_mV_ | pDE43-MCS Strepto^R^ | In-fusion cloning | PCR using pGMCS-*P_pacL1_*-*pacL1*_mT_-*ctpC*_mV_ as template with Inf-3269Del(53-86)-left/Inf-3269C7-right primers and circularization by In-fusion reaction | This work |
| pGMCS-*P_pacL1_*-*pacL1*(E^55^V)-*ctpC* | pDE43-MCS Strepto^R^ | In-fusion cloning | PCR using pGMCS-*P_pacL1_*-*pacL1*-*ctpC* as template with subs EA 55-59 Left/subs EA 55 Right primers and circularization by In-fusion reaction | This work |
| pGMCS-*P_pacL1_*-*pacL1*(E^59^A)-*ctpC* | pDE43-MCS Strepto^R^ | In-fusion cloning | PCR using pGMCS-*P_pacL1_*-*pacL1*-*ctpC* as template with subs EA 59 Left/subs EA 59 Right primers and circularization by In-fusion reaction | This work |
| pGMCS-*P_pacL1_*-*pacL1*(E^71^A)-*ctpC* | pDE43-MCS Strepto^R^ | In-fusion cloning | PCR using pGMCS-*P_pacL1_*-*pacL1*-*ctpC* as template with Rv3269(E71A) subst Left/Rv3269(E71A) subst Right primers and circularization by In-fusion reaction | This work |
| pGMCS-*P_pacL1_*-*pacL1*(3EA)-*ctpC* | pDE43-MCS Strepto^R^ | In-fusion cloning | PCR using pGMCS-*P_pacL1_*-*pacL1(E^71^A)*-*ctpC* as template with subs EA 55-59 Left/subs EA 55-59 Right primers and circularization by In-fusion reaction | This work |
| pGMCS-*P_pacL1_*-*pacL1*(E^71^A)_Flag_-*ctpC*_His6_ | pDE43-MCS Strepto^R^ | In-fusion cloning | PCR using pGMCS-*P_pacL1_*-*pacL1*_Flag_-*ctpC*_His6_ as template with Rv3269(E71A) subst Left/Rv3269(E71A) subst Right primers and circularization by In-fusion reaction | This work |
| pGMCS-*P_pacL1_*-*pacL1*(3EA)_Flag_-*ctpC*_His6_ | pDE43-MCS Strepto^R^ | In-fusion cloning | PCR using pGMCS-*P_pacL1_*-*pacL1*(E71A) _Flag_-*ctpC*_His6_ as template with subs EA 55-59 Left/subs EA 55-59 Right primers and circularization by In-fusion reaction | This work |
| pGMCS-*P_pacL1_*-*pacL1*(3EA)_mT_-*ctpC*_mV_ | pDE43-MCS Strepto^R^ | In-fusion cloning | PCR using pGMCS-*P_pacL1_*-*pacL1*(E71A) _mT_-*ctpC*_mV_ as template with subs EA 55-59 Left/subs EA 55-59 Right primers and circularization by In-fusion reaction | This work |
| pGMCS-*P_pacL1_*-*pacL2* | pDE43-MCS Strepto^R^ | In-fusion cloning | *pacL2* fragment amplified with H37Rv chromosomal DNA and Inf-1993c-Am/Inf-1993c-Av cloned by In-fusion reaction in the backbone amplified from pGMCS-*P_pacL1_*-*pacL1* with Inf-Rv3269-left/Inf-Rv3269-right | This work |
| pGMCS-*P_pacL1_*-*pacL3* | pDE43-MCS Strepto^R^ | In-fusion cloning | *pacL3* fragment amplified with H37Rv chromosomal DNA and Inf-0968-Am/Inf-0968-Av cloned by In-fusion reaction in the backbone amplified from pGMCS-*P_pacL1_*-*pacL1* with Inf-Rv3269-left/Inf-Rv3269-right | This work |
| pGMCS-*P_pacL1_*-*pacL2*-*ctpC* | pDE43-MCS Strepto^R^ | In-fusion cloning | *pacL2* fragment amplified with H37Rv chromosomal DNA and Inf-1993-Am/Inf-1993-Av cloned by In-fusion reaction in the backbone amplified from pGMCS-*P_pacL1_*-*pacL1*-*ctpC* with Inf-Rv3269-left/Inf-Rv3269-right | This work |
| pGMCS-*P_pacL1_*-*pacL2*(MBM)-*ctpC* | pDE43-MCS Strepto^R^ | In-fusion cloning | *pacL2*(MBM) fragment amplified with H37Rv chromosomal DNA and Inf-1993-Am/Inf-1993MBM-Av cloned by In-fusion reaction in the backbone amplified from pGMCS-*P_pacL1_*-*pacL1*-*ctpC* with Inf-Rv3269-left/Inf-Rv3269-right | This work |
| pGMCS-*P_pacL1_*-*pacL3*-*ctpC* | pDE43-MCS Strepto^R^ | In-fusion cloning | *pacL3* fragment amplified with H37Rv chromosomal DNA and Inf-0968-Am/Inf-0968-Av cloned by In-fusion reaction in the backbone amplified from pGMCS-*P_pacL1_*-*pacL1*-*ctpC* with Inf-Rv3269-left/Inf-Rv3269-right | This work |
| pGMCS-*P_pacL1_*-*pacL1*_mT_-*pacL2*_mV_ | pDE43-MCS Strepto^R^ | In-fusion cloning | *pacL2* fragment amplified with H37Rv chromosomal DNA and Rv1993c-XFP-Fw/ Rv1993c-XFP-Rv cloned by In-fusion reaction in the backbone amplified from pGMCS-*P_pacL1_*-*pacL1*_mT_-*ctpC*_mV_ with mT av left/infus backbone mVenus Right primers | This work |
| pGMCS-*P_pacL1_*-*pacL1*_mT_-*pacL3*_mV_ | pDE43-MCS Strepto^R^ | In-fusion cloning | *pacL3* fragment amplified with H37Rv chromosomal DNA and Rv0968-XFP-Fw/ Rv0968-XFP-Rv cloned by In-fusion reaction in the backbone amplified from pGMCS-*P_pacL1_*-*pacL1*_mT_-*ctpC*_mV_ with mT av left/infus backbone mVenus Right | This work |
| pGMCS-TetR-P1-MbcAT | pDE43-MCS StreptoR | - | - | ^3^ |
| pGMCS-TetR-P1-*pacL1*_Flag_-*P_pacL1_*-*ctpC*_His6_ | pDE43-MCS StreptoR | In-fusion cloning | A TetR-encoding DNA fragment was amplified using pGMCS-TetR-P1-MbcAT as template and TetR-Am-Fus-P1/TetR-Av-Rv primers. A P1 promoter fragment was amplified using pGMCS-TetR-P1-MbcAT as template and Inf-P1-left/Tet-P1-Fus-Fw primers. A PacL1_Flag_-encoding fragment was amplified using pGMCS-*P_pacL1_*-*pacL1*_Flag_-*ctpC*_His6_ as template with Rv3269-P1-Fus-Fw/Rv3269-Flag-Rev primers. These fragments were cloned by In-Fusion reaction in the backbone amplified from pGMCS-*PpacL1*-*pacL1*_mT_-*ctpC*_mV_ with ctpC-B2-left/ctpC-B2-right primers. | This work |
| pProEx-Htb | Ampi^R^ |  | Expression vector with IPTG-inducible trc promoter, His6 tag and TEV cleavage site. | Invitrogen |
| pProEx-Htb-SolPacL1 | pProEx-Htb; Ampi^R^ | Restriction site cloning | A DNA fragment encoding SolPacL1 was amplified using H375v chromosomal DNA and SolPacL1Fw/SolPacL1Rv primers, cut with *NcoI* and *HindIII* and ligated in pProEx-Htb cut with *NcoI* and *HindIII* | This work |
| pProEx-Htb-SolPacL1Δ3 | pProEx-Htb; Ampi^R^ | Restriction site cloning | pProEx-Htb-SolPacL1 DNA was amplified with fragment encoding SolPacL1 was amplified with SolPacL1Δ3Fw/SolPacL1Δ3Rv primers, cut with *HindIII* and self-ligated | This work |

Table S3. Related to Materials and Methods; Primers used for plasmid constructions.

| **Name** | **Sequence 5'-> 3'** |
| --- | --- |
| 3269 Cter Flag-Am-ctpC-Right | GCACGACCACGACCACGACTACAAGGACGACGA  TGACAAATGAGCGCCTCGCCATGACC |
| ctpC Cter His-Av-attB3-Right | TACCGCCTGGACCGCCACCACCACCACCACCACT  GAGATAATTCACTGGCCGTCG |
| clo-B2-3269-Am | GGGGACAGCTTTCTTGTACAAAGTGGAGCGATACT  CGACGATTC |
| clo-B3-3269-Av | GGGGACAACTTTGTATAATAAAGTTGGCTCAGTGGT  CGTGGTCGTGC |
| clo-B3-ctpC-Av | GGGGACAACTTTGTATAATAAAGTTGGCTAGCGGTC  CAGGCGGTAGCGG |
| Del3269-Cterm3-Right | ACTCCAGCGATCAGCGACCTGCACGACTGAGCCA  ACTTTATTATACATAGTTGATAATTCACTGG |
| Del3269-Cterm7-Right | ACTCCAGCGATCAGCTGAGCCAACTTTATTATAC  ATAGTTGATAATTCACTGG |
| Del3269-Cterm-Left | GCTGATCGCTGGAGTGGGCG |
| Del3269-Nterm7_26-Right | GCGATACAAGTGTTCTTAAAAAAGGCCGCGGCCAAAG |
| Del3269-Nterm-Left | GAACACTTGTATCGCCATTGCCTG |
| Inf-0968-Am | AAGGTACAGGCAATGGTGTGGCATGGATTCCTAG |
| Inf-0968-Av | TGGTCGTGGTCGTGCTCAGTGGTCATGACCGTCG |
| Inf-1993-Am | AAGGTACAGGCAATGGTTACGCATGAGCTATTGG |
| Inf-1993-Av | TGGTCGTGGTCGTGCTCACTCGTCGACCCTGGCGCCAG |
| Inf-1993MBM-Av | TGGTCGTGGTCGTGCAGGTCGGCGCCAGCGGGCAGGGGCG |
| Inf-3269C7-right | GACCTGCACGACCACGACCAC |
| Inf-3269Del(37-86)-left | GTGGTCGTGCAGGTCAAGCGGCGCTTTGGCCGCGG |
| Inf-3269Del(53-86)-left | GTGGTCGTGCAGGTCCTTGCGGGTTCCGCGCAGACC |
| Inf-3269DelC7-left | GTCGTCCTTGTAGTCGCTGATCGCTGGAGTGGGCG |
| InF-ctpC-MutPB-L1 | CTCGTTGCAGTGACATTAAGGTCTATATATCGCGAT  ATCAATATG |
| InF-ctpC-MutPB-R1 | TGTCACTGCAACGAGCTGCCG |
| Inf-Flag-right2 | GACTACAAGGACGACGATGAC |
| Inf-P1-left | GAAATGATGTATGCCGTGCTGGTC |
| Inf-Rv3269-left | CATTGCCTGTACCTTTCTTCC |
| Inf-Rv3269-right | GCACGACCACGACCACTGAG |
| infus backbone mVenus Right | CTGGAGGGTTCGGGCGTGAG |
| infus mCherry Rv | TTTACCTTCCTCGCCTTACTTGTACAGCTCGTCCA |
| infus mVenus Rv1488 Rv | CACGCCCGAACCCTCCAGTTGAGTCAACCTGGGGGGC |
| infus P1 Rv1488 Fw | ATGAGAGGAGGATTCAC |
| Infus pEN41A-P1_x_mCherry Right | GGTGAATCCTCCTCTCATT |
| infus P1 Rv1488 Fw (pGMC) | AGAGCCTGCGGCATGCATGCAAGGAGCCGTTGCT |
| infus Rv1488 Fw | CAAGTAAGCGCCTCGCCGTGCAAGGAGCCGTTGCT |
| mCherry Fw2 w/o start codon | GTGAGCAAGGGCGAGGA |
| mT av left | GGCGAGGCGCTTACTTGTAG |
| mTurquoise-Fw #3 | CACGACCACGACCACCTGGAGGGTTCGGGCGTCAGC  AAGGGTGAGGAAC |
| mTurquoise-Rv #3 | GTCATGGCGAGGCGCTTACTTGTAGAGTTCGTCCATGC |
| mVenus-Fw #1 | GCTACCGCCTGGACCGCCTgGAGGGTTCgGGcGTGAGC  AAGGGCGAGGAGC |
| mVenus-Rv #1 | CGGCCAGTGAATTATCGGGGTTACTTGTACAGCTCGT  CCATGCCG |
| pGMC-12-left | GGCGAGGAAGGTAAACAGG |
| pGMC-12-right | CATGCCGCAGGCTCTCTTTG |
| pGMC 3269 Am Left | CATTGCCTGTACCTTTCTTCC |
| pGMC-3269-FLUO-ctpC Left | TCCTCGCCCTTGCTCACCCCCGAACCCT |
| pGMC-Am-ctpC Right #3 | GCGCCTCGCCATGACC |
| pGMC-Av-attB3-Right #1 | GATAATTCACTGGCCGTCG |
| pGMC del3269 Right | AAGGTACAGGCAATGACCCTGGAAGTGGTATCG |
| pGMC-Cter-ctpC Left #1 | GCGGTCCAGGCGGTAGCG |
| pGMC-Cter-rv3269 Left #3 | GTGGTCGTGGTCGTGC |
| Rv0968 XFP Fw | CAAGTAAGCGCCTCGCCAGGAGCATCCGCGATGGT |
| Rv0968 XFP Rv | TCCTCGCCCTTGCTCACGCCCGAACCCTCCAGGTGGT  CATGACCGTCGTCCG |
| Rv1993c XFP Fw | CAAGTAAGCGCCTCGCCAGGAGAACTGACCGTGGT |
| Rv1993c XFP Rv | CTCCTCGCCCTTGCTCACGCCCGAACCCTCCAGCTCG  TCGACCCTGGCGCC |
| Rv3269(E71A) subst Left | GGCCATCACGTCGGCCA |
| Rv3269(E71A) subst Right | CCGACGTGATGGCCGCCGCTCGTGAGCGCATC |
| Rv3269-Flag-Rev | ATCGGCGGGAGAATCGCGCTCATTTGTCATCGTCG |
| Rv3269-mT-Rev | ATCGGCGGGAGAATCGGCGAGGCGCTTACTTGTAG |
| Rv3269-P1-Fus-Fw | CAGCACGGCATACATCATTTCGGAAGAAAGGTACAGGCAATGG |
| SolPacL1Δ3-Fw | GACCTGCACGACTAGGACCACTGAAAGCTTGGC |
| SolPacL1Δ3-Rv | GCCAAGCTTTCAGTGGTCCTAGTCGTGCAGGTC |
| SolPacL1Rv | CCAGGGTCATGGCGAAGCTTTCAGTGGTCGTGGTCGTGC |
| SolPacL1Fw | CCGCCTACGAGATCTTAACCATGGCCGCGGCCAAAGCG |
| subs EA 55 Right | GAACCCGCAAGGCCGTGGAAGCCGCGGAATCGG |
| subs EA 55-59 Left | CGGCCTTGCGGGTTCCGCG |
| subs EA 55-59 Right | GAACCCGCAAGGCCGTGGAAGCCGCGGccTCGGCCCG  CCTAAAGG |
| subs EA 59 Left | CCGCGGCTTCCTCGGCCTTG |
| subs EA 59 Right | CCGAGGAAGCCGCGGCCTCGGCCCGCCTAAAGG |
| Tet-P1-Fus-Fw | CGATCCAATATTACGAGATCGGATCGTCGGCACCGTCAC |
| TetR-Am-Fus-P1 | GATCTCGTAATATTGGATCG |
| TetR-Av-Rv | GGAGCGATACTCGACGTTCTCGGCTCGATGATCCC |

Table S4. Related to Materials and Methods; Primers used for RT-qPCR experiments.

| **Gene** | **Name** | **Sequence 5'-> 3'** |
| --- | --- | --- |
| *pacL1* | RT 3269 Fw | TGGCGATACAAGTGTTCTTGGCG |
|  | RT 3269 Rv | ACGTCGGCCACCTTTAGGCGG |
| *pacL1 (N-term)* | RT 3269Nt Fw1 | CAAGTGTTCTTGGCGAAGG |
|  | RT 3269Nt Rv1 | TAAGATCTCGTAGGCGGTCA |
| *ctpC* | RT ctpC Fw | TCACCATTTTCACCGGGTAT |
|  | RT ctpC Rv | GATGTTGAGCAACCACAGGA |
| *sigA* | RT sigA_2F | AAGACACCGACCTGGAACTC |
|  | RT sigA_2R | CGGCATCAGCTTCTTCTTC |
| *rpoB* | RT rpoB Fw | TCGTTCTCTGACCCTCGTTTC |
|  | RT rpoB Rv | ACGTGCCCTTCTCGGTCATCA |
